# A prospective longitudinal study shows putamen volume is associated with moderate amphetamine use and resultant cognitive impairments

**DOI:** 10.1101/2020.10.29.361378

**Authors:** Keith M Kendrick, Joerg Daumann, Daniel Wagner, Philip Koester, Marc Tittgemeyer, Qiang Luo, Euphrosyne Gouzoulis-Mayfrank, Benjamin Becker

**Affiliations:** Key Laboratory for Neuroinformation, Center for Information in Medicine, School of Life Science and Technology, University of Electronic Science and Technology of China, Chengdu, China; Department of Psychiatry and Psychotherapy, University of Cologne, Germany; Max-Planck Institute for Neurological Research, Cologne, Germany; Institute of Science and Technology for Brain-Inspired Intelligence, Ministry of Education Key Laboratory of Computational Neuroscience and Brain-Inspired Intelligence, Fudan University, Shanghai, 200433, PR China; LVR Clinics of Cologne, Cologne, Germany

**Author notes:** Correspondence to: Benjamin Becker, University of Electronic Science and Technology of China, Center for Information in Medicine, No. 2006, Xiyuan Ave, West Hi-Tech Zone, 611731, Chengdu, China. Contributed equally to this work (joint first authors) (Keith M Kendrick, Joerg Daumann).

**Keywords:** Amphetamines, brain volume, dorsal striatum, MDMA, prospective, stimulants

## Abstract

**Background:** Amphetamine-type stimulants (ATS) have become a critical public health issue. Animal models have indicated a clear neurotoxic potential of ATSs. In humans, chronic use has been associated with cognitive deficits and structural brain abnormalities. However, cross-sectional retrospective designs in chronic users cannot truly determine the causal direction of the effects.

**Methods:** In a prospective-longitudinal study design cognitive functioning and brain structure were assessed at baseline and at 12-months follow-up in occasional ATS users (cumulative lifetime use <10 units at baseline).

**Results:** Examination of change-scores between the initial examination and follow-up revealed declined verbal memory performance and putamen volume in users with high relative to low interim ATS exposure. In the entire sample interim ATS use, memory decline and putamen volume reductions were strongly associated.

**Conclusions:** The present findings support the hypothesis that ATS use is associated with deficient dorsal striatal morphology which might reflect alterations in dopaminergic pathways. More importantly, these findings strongly suggest that even occasional, low-dose ATS use disrupts striatal integrity and cognitive functioning.

## Introduction

Increasing rates of recreational amphetamine-type stimulant (ATS) use, predominately illicitly produced amphetamine (AMPH) and 3,4-methylenedioxymethamphetamine (MDMA, ‘Ecstasy’) and of ATS users seeking treatment indicate that ATSs have become a major health problem (UNODC, 2011; 2014). In terms of prevalence rates ATS is second only to cannabis (UNODC, 2011), with recreational use among, often socially-well integrated, young adults being the most typical pattern (Gouzoulis-Mayfrank et al., 2009). During the last decades converging evidence from different animal models indicates a neurotoxic potential of ATSs (Aguilar et al., 2020; Parrott, 2013; Moratalla et al., 2017).These animal studies have shown that the experimental application of varying dosage regimens of MDMA and amphetamines lead to long-term neurotoxic effects in rodent and nonhuman primate models, as indicated by a range of brain morphological and neurochemical indices (overview see e.g. Moratalla et al., 2017). However, the key question as to whether human ATS users may suffer from similar neurotoxic brain lesions remains unanswered.

Convergent evidence from animal models and meta-analyses covering neuroimaging studies in human drug users suggest that prolonged drug use is associated with structural and functional adaptations in limbic-striato-prefrontal circuits of the brain (Everitt and Robbins, 2016; Klugah-Brown et al., 2020; Ersche et al., 2013). Accumulating evidence from human studies suggests that the chronic use of ATS is associated with altered brain morphology, particularly deficient grey matter (GM) integrity in limbic-striato-prefrontal brain networks, as well as subtle yet consistently observed, deficits in cognitive and emotional functions that have been associated with this circuitry (Gouzoulis-Mayfrank et al., 2009; Wagner et al., 2013; Ersche et al., 2013; Mackey & Paulus, 2013; Parrott 2015). However, the majority of human findings are based on cross-sectional studies in the sub-group of chronic, often dependent, users of the more-addictive amphetamine compound methamphetamine (MA), also known as ‘Crystal-meth’. Due the retrospective design and the lack of baseline data these studies do not allow a separation of specific effects of ATS use, such as potential neurotoxic effects or addiction-related brain-plastic adaptations, from alterations that precede, or promote, the onset of use. Only longitudinal designs that control for baseline differences can truly determine whether the neuropsychological or neuroanatomical differences in ATS users are a result of drug use or a predisposing factor (Taylor et al., 2013).

Using sophisticated sampling strategies in cross-sectional study designs, that also include more appropriate control groups and prospective designs, we and others have begun to disentangle the contribution of predisposing and drug-associated factors in brain structural abnormalities observed in ATS users (Daumann et al., 2011; Ersche et al., 2012; Becker et al., 2015). Findings from these studies suggest that GM alterations in regions associated with emotional and cognitive control, particularly the amygdala, the anterior cingulate and adjacent medial prefrontal regions prior to the onset of ATS use may represent reliable brain-structural vulnerability markers for increased risk to develop escalating use and potential addiction. However, studies with longitudinal-designs specifically focusing on brain-structural effects of ATS users while controlling for baseline abnormalities are rare.

The assessment of brain structural changes in longitudinal designs has additionally been hampered by methodological issues. Traditional longitudinal voxel-based morphometry (VBM, Ashburner & Friston, 2000) analyses use simple intra-subject registration approaches and asymmetric processing that bias the estimation of longitudinal changes (Ashburner & Friston, 2011; Thompson & Holland, 2011). More recent developments in longitudinal VBM techniques, such as group-wise intra-subject models (symmetric approaches) that combine rigid-body and diffeomorphic (Ashburner & Friston, 2011) registration and correction for inhomogeneity artefacts (Ashburner & Ridgway, 2012) have enabled researchers to evaluate brain structural changes with more appropriate statistical models and accordingly a higher sensitivity for longitudinal changes.

Against this background we applied the optimized VBM machinery to a longitudinal brain structural dataset acquired in occasional ATS users with only minimal ATS exposure at study inclusion (cumulative lifetime use < 10 units of ATS) to specifically examine the long-term effects of ATS use on brain structure while controlling for baseline differences and other known confounders in this field (e.g. co-use of other drugs, particularly cannabis (Gouzoulis-Mayfrank & Daumann, 2006). To this end brain structure, cognitive functioning and interim drug use were re-assessed after a follow-up period of 12-months. Using a data-driven clustering approach users with low **(LOW)** and high **(HIGH)** ATS use during follow-up were identified. Next, cognitive domains and brain regions with differential between-group changes during follow-up were explored using a correlational approach to take advantage of the entire sample of n = 17 in examining ATS-use associated functional and structural changes.

## Materials and methods

### Participants

Participants in the present study were a sub-group of a larger research project and their baseline data had been used for cross-sectional brain-structural comparisons (Daumann et al., 2011; Becker et al., 2015). The main inclusion criterion at baseline was occasional (ATS use > 1 occasion), but very limited use of ATS (cumulative lifetime use of <10 units of ATS). In line with previous studies (Daumann et al., 2011; Becker et al. 2015) units were defined on the basis of typical quantities that the MDMA and amphetamine are supplied in (one unit MDMA = 1 tablet; one unit amphetamine = 1 g). In addition the following exclusion criteria were used: lifetime use of any other illicit psychotropic substances > 5 occasions (except for cannabis, which is widely used among recreational ATS users), history of alcohol abuse or dependence (according to DSM-IV criteria), regular medication (once or more a week, except for contraceptives), use of any psychotropic substances in the 7 days before the examination (exception: cannabis, tobacco), use of cannabis on the day of the examination, current or previous history of neurological or psychiatric disorder (Axis I and II according to DSM-IV criteria), any other general medical condition, history of traumatic brain injury with loss of consciousness or amnesia, left-handedness, unable to give informed consent, age at least 18 years, childhood diagnosis of attention-deficit hyperactivity disorder, pregnancy and MRI contraindications. Importantly, a previously published cross-sectional comparison with drug-naïve subjects revealed no brain-structural alterations in the group of occasional ATS users (Daumann et al., 2011). In addition, cognitive functioning as well as a range of potential confounders, including use of other licit and illicit drugs, psychopathology, cognitive functioning and urine as well as hair samples to validate drug use patterns were assessed (details see Wagner et al., 2013; Becker et al., 2013). Follow-up brain structural data could be assessed in n = 19 from the n = 42 participants that were included during the baseline assessments. Cognitive performance was assessed using a comprehensive, validated neurocognitive test battery including measures of verbal and visual long- and short-term memory, speed of information processing, cognitive inference and flexibility ((20)). After detailed study description subjects provided written informed consent; the study had full ethical approval by the Medical Faculty of the University of Cologne and was in accordance with the latest revision of the Declaration of Helsinki.

### Procedures

At baseline, 42 occasional ATS users were enrolled in the cross-sectional study (for details see Daumann et al., 2011). After baseline assessment of brain structure, drug use, cognitive performance and potential confounders participants were followed to re-assess brain structure, cognitive functioning and interim ATS use during a 12-months follow-up interval. At follow-up brain structure could be re-assessed in a total of n = 19 participants. Screening procedures included a structured interview to assess Diagnostic and Statistical Manual of Mental Disorders-Fourth edition (DSM-IV) Axis I and II disorders, the Wender Utah Rating Scale (Ward et al., 1993) to assess childhood attention-deficit hyperactivity disorder, a detailed structured drug-history interview for ATS and other prevalent psychotropic substances. Randomly taken hair samples and urine screens were used to verify self-reported substance use patterns. In addition, the following potential confounding variables were assessed: neuropsychological functioning, including memory, executive functioning, mental flexibility, non-verbal intelligence, use of alcohol and tobacco, overall psychological distress (Global Severity Index from the Symptom Checklist-90-R, SCL90R).

### Cognitive test battery

#### Auditiv-Verbaler Lerntest AVLT

Verbal declarative memory performance was examined by the German version (Auditiv-Verbaler Lerntest (AVLT) (Heubrock, 1992) of the Rey Auditory Verbal Learning Test (Rey, 1964). The RAVLT assesses verbal declarative memory performance by means measures of immediate recall, total acquisition performance across five trials, recall after interference, loss after interference and recognition after 30 minutes.

#### Lern- und Gedächtnistest LGT 3

Visual paired associates learning was assessed by a subtest of The Lern- und Gedächtnistest (LGT) (Baeumler, 1974). The sub-test contains figures composed of a logo and surrounding frame presented to the subject for 60 seconds. Subjects have to choose the correct logo-frame composition from 4 options, immediately after the presentation (immediate recall) and after a delay of 1 hour (delayed recall).

#### Digit-Span-Backward

Is a classical working memory measure from the Hamburg-Wechsler-Intelligenztest für Erwachsene (HAWIE-R) (Tewes, 1991), a German version of the Wechsler Intelligence Test (WAIS) (Wechsler, 2008). Subjects listen to a sequence of digits and have to recall the digits immediately in reverse order.

#### Digit symbol test

This test from the WAIS (Wechsler, 2008) (German Version HAWIE-R) assesses speed of information processing using a total of nine digit-symbol pairs (e.g. 1/-,2/_…7/L,8/X,9/=) followed by a list of 93 digits. Subjects are required to write down the corresponding symbol for each digit as quickly as possible. Number of correct symbols within 90 seconds is the most commonly used measure of performance.

#### Stroop task

A classical stroop task (German Version, Farbe-Wort-Interferenztest, (Stroop 1935, Baeumler, 1985)) assesses cognitive interference/inhibition. Speed of performance, corrected errors and uncorrected responses are assessed for reading color-names, color rectangles and color-names in different color inks (inference condition).

#### Trail-making test

This classical test of mental flexibility (Raitan, 1992) requires subjects to connect circles numbered from 1 to 25 (Part A) and numbers (1-13) and letters (A-L) alternatively (Part B). Response times are recorded.

#### Raven Standard Progressive Matrices

Baseline differences in general intelligence at baseline were assessed using the Raven Standard Progressive Matrices (Raven et al., 1998).

### MRI data acquisition and analysis approach

High-resolution brain structural MRI data was acquired on a 3 Tesla Magnetom Tim Trio system using a standard quadrature head coil (flip angle = 18°, repetition time = 1930 ms, echo time = 5.8 ms, slice thickness = 1.25 mm, voxel size = 1.0 × 1.0 × 1.25 mm). Analyses of the longitudinal data were carried out using optimized segmentation protocols and the new longitudinal registration module in SPM12 (18). In line with previous studies (20, 24) ATS-use associated changes in cognitive functioning were assessed by means of change scores between baseline and follow-up. In accordance with this approach, effects on GM volume were assessed using individual differential GM maps (baseline vs. follow-up). Groups were directly compared using independent t-tests. To increase the sensitivity to detect brain structural changes the analyses focused on key brain structures associated with ATS use (Ersche et al., 2013; Mackey & Paulus, 2013; Becker et al., 2015 namely basal ganglia, amygdala, medial prefrontal cortex, inferior frontal gyrus, and insula, using structural regions of interest (ROIs). Structural regions of interest were defined using the Anatomy Toolbox version 1.8 (Eickhoff et al., 2005) and the WFU Pickatlas Toolbox (Maldjian et al., 2003). Between-group differences within the a priori regions of interest were computed using a threshold of P < .05 (family-wise error-corrected, FWE). Results were thresholded at a family-wise error corrected (FWE) p < 0.05. For the analyses, variables that were not normally distributed, including interim ATS use, were initially log-transformed to achieve a normal distribution.

## Results

Based on automatized standard quality assessments of MRI data one subject was excluded from all further analyses. Participants had used a mean of 7.72 (SD 8.99, range 0-27) units of ATS during the follow-up period. One user reported having used 74.2 units of ATS during follow-up and was excluded as outlier from all further analyses (z = 3.51). Based on the reported log-transformed ATS use during follow-up data-driven k-means clustering with squared Euclidean distance revealed two separate sub-groups of users with low (**LOW**, n = 11) and high ATS use (**HIGH**, n = 8). Users in the **LOW**(n = 10) group had used a mean of 1.45 (SD 1.27, range 0-3.50) units of ATS, whereas those in the **HIGH**(n = 7) group had used a mean of 16.69 (SD 7.33, range 8.80-18.20) units during the follow-up (paired t-test, t = −6.52, df = 15, p < 0.001). Importantly, groups did not show differences on a range of potential confounders at baseline, including socio-demographics and pre-baseline drug use compared to follow-up, including days between the scanning sessions and interim cannabis use (**table 1**).

**Table 1:** Demographic and drug-use characteristics of the groups.

|  | **LOW (n=10)** | **HIGH (n=7)** | **p-value** |
| --- | --- | --- | --- |
| **At study inclusion** |  |  |  |
| Age | 24.50 ($\pm 5.38$) | 22.29 ($\pm 5.52$) | 0.42 |
| Education (years) | 15.45 ($\pm 2.92$) | 13.32 ($\pm 2.56$) | 0.14 |
| General intelligence (Raven, errors) | 6.40 ($\pm 4.35$) | 7.57 ($\pm 8.67$) | 0.72 |
| No of cigarettes / day | 6.30 ($\pm 6.83$) | 10.07 ($\pm 7.07$) | 0.28 |
| Years of tobacco use | 4.95 ($\pm 6.08$) | 5.14 ($\pm 4.22$) | 0.94 |
| No alcohol drinks / week | 8.00 ($\pm 1.82$) | 8.28 ($\pm 0.76$) | 0.71 |
| Age cannabis use onset | 15.30 ($\pm 2.41$) | 16.29 ($\pm 6.62$) | 0.67 |
| Frequency of cannabis use (days/month) | 16.15 ($\pm 11.31$) | 16.14 ($\pm 14.61$) | 0.99 |
| ATS cumulative (units) | 5.10 ($\pm 2.60$) | 6.10 ($\pm 1.88$) | 0.40 |
| **During follow-up** |  |  |  |
| Days between t1 – t2 | 447.50 ($\pm 109.95$) | 382.28 ($\pm 54.13$) | 0.17 |
| ATS cumulative (units) | 1.45 ($\pm 1.27$) | 16.68 ($\pm 7.33$) | <0.001** |
| No of cigarettes / day | 7.40 ($\pm 9.03$) | 7.29 ($\pm 7.95$) | 0.97 |
| No alcohol drinks / week | 8.60 ($\pm 1.35$) | 7.00 ($\pm 1.83$) | 0.06 |
| Frequency of cannabis use (days/month) | 15.15 ($\pm 11.41$) | 16.00 ($\pm 10.96$) | 0.88 |
| Cannabis cumulative (gram) | 84.36 ($\pm 83.51$) | 125.14 ($\pm 111.8$) | 0.40 |

Analyses of change scores revealed significant differences between the **HIGH** and **LOW** groups only in the domain of verbal memory (total number of words recalled across 5 trials of a word list; Rey Auditory Verbal Learning Test, RAVLT, Rey 1964) (t = 2.347, df = 15, p = 0.032). Compared to the baseline assessment users in the **LOW** group remembered on average 2.1 (SD = 4.5) words more at follow-up, whereas the **HIGH** group remembered on average 4.0 (SD = 6.13) words less at follow-up (**Fig. 1a**). The groups did not differ on change scores for other cognitive measures (all p > 0.07). Analyses of brain structural data revealed a significant interaction effect in the basal ganglia located in the right putamen (t = 4.31, p < 0.05, maximum at 30 / 8 / −9 **Fig. 2a**). Extraction of individual GM volumes from this region further revealed that this effect was driven by a significant reduction in the **HIGH** group (t = 4.07, df = 6, p = 0.007), whereas GM indices did not change significantly in the **LOW** group (p = 0.148) (**Fig. 1b**). A correlational analysis that took advantage of the entire sample revealed a significant negative association between interim ATS use and GM changes (n = 17, r = − 0.72, R^2^ = 0.51, p = 0.001), indicating a direct association between the amount of interim ATS use and GM reductions in the right putamen. In addition, the change in the total number of words remembered in the RAVLT (follow-up minus baseline) significantly correlated with both GM changes in the right putamen (n = 17, r = 0.53, R^2^ = 0.28, p = 0.029) as well as the amount of interim ATS use (n = 17, r = −0.59, R^2^ = 0.35, p = 0.012), indicating that a higher loss of words recalled was associated with higher putamen decreases as well as higher ATS use during follow-up (correlations are shown in **Fig. 2b**). Moreover, higher differences in the RAVLT immediate recall during the first learning trial were trend-to-significant related to higher interim ATS use (n = 17, r = −0.44, p = 0.075).

**Figure 1:**
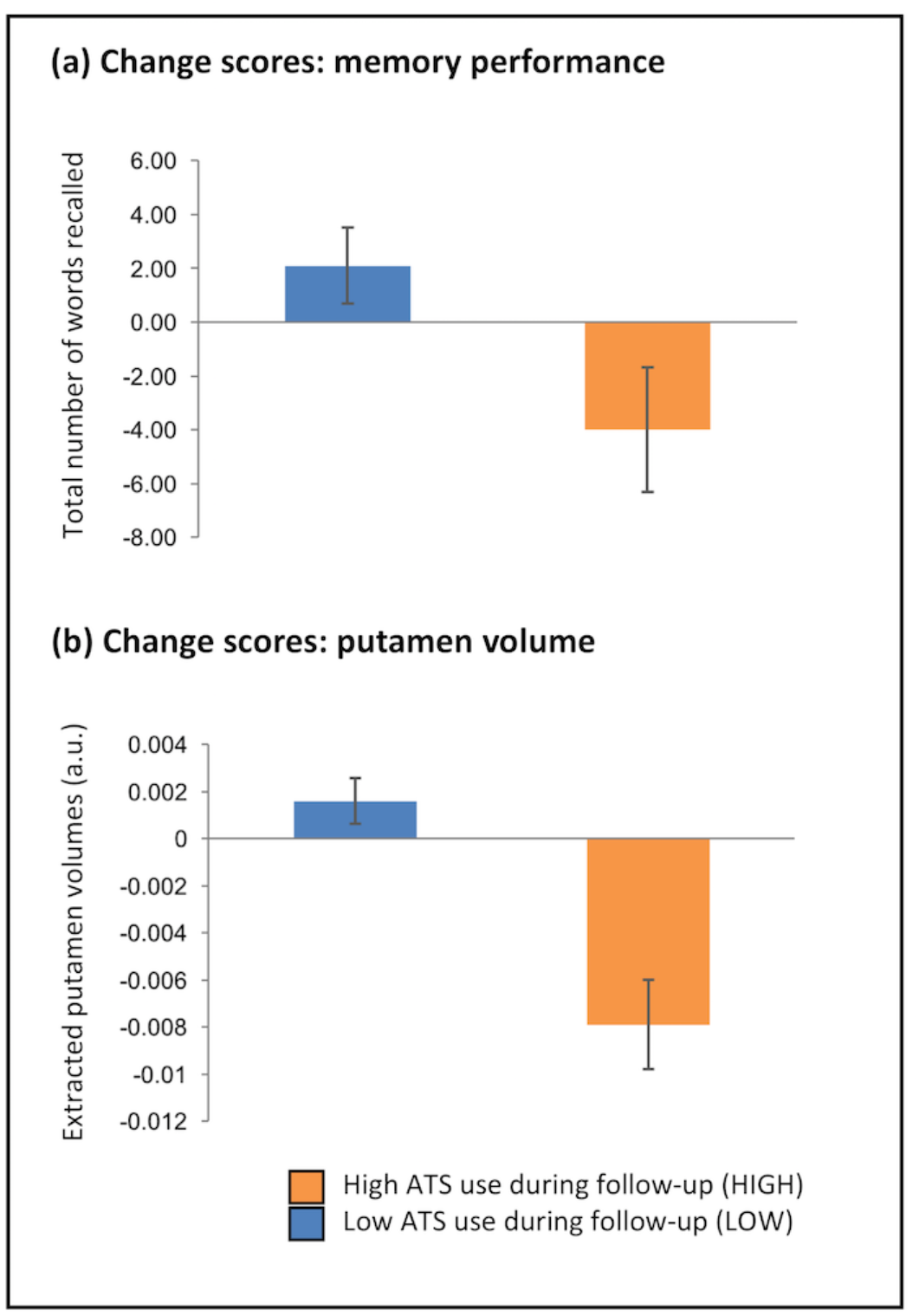
Memory performance and putamen GM change-scores for the groups. Change scores (baseline vs follow-up) from verbal memory performance (a) and putamen GM (b). Users with higher ATS-use (HIGH) during follow-up demonstrated significant performance and putamen GM loss relative to users with low ATS-use (LOW) during follow-up.

**Figure 2:**
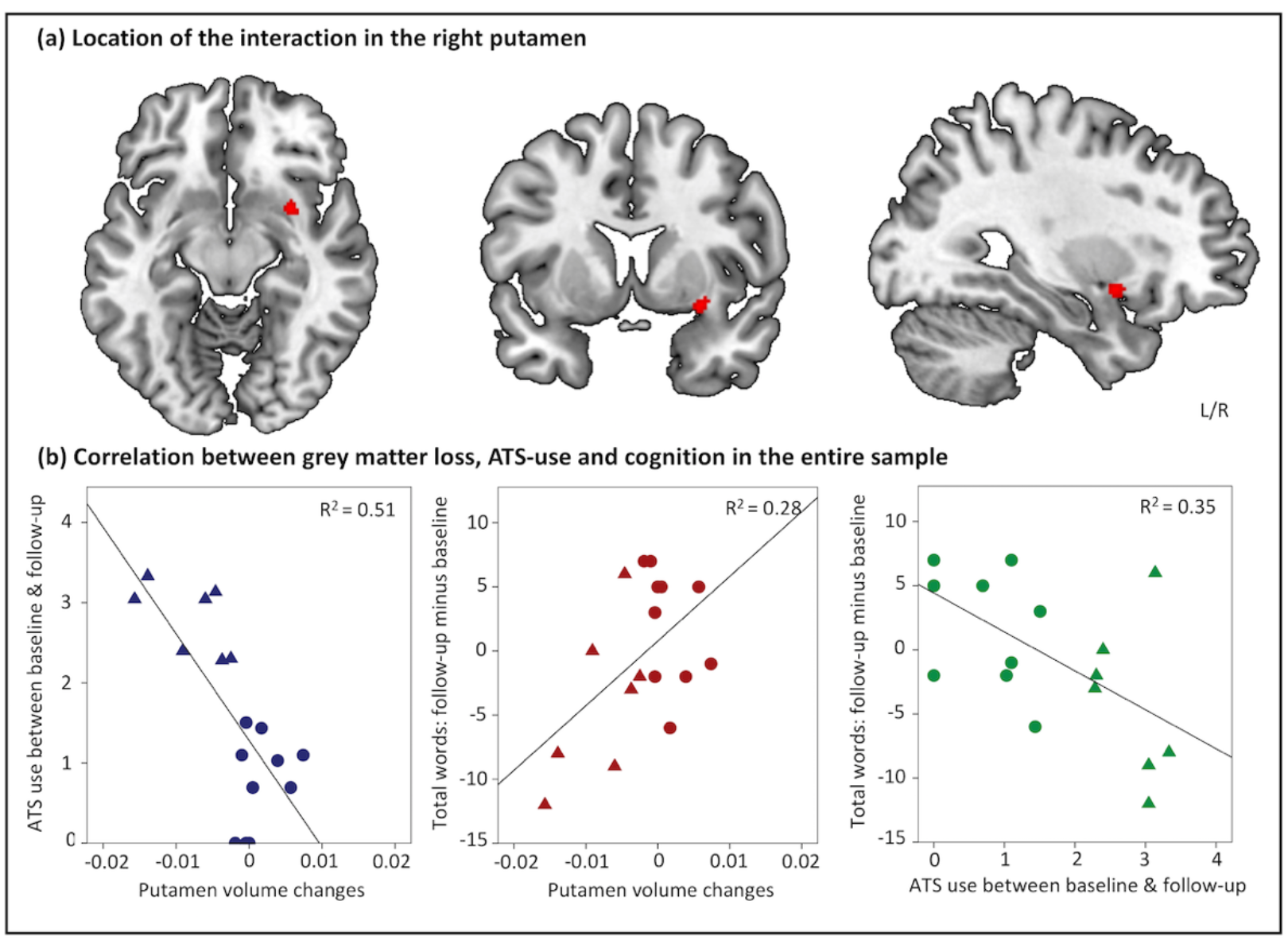
Comparison of gray matter and associations with ATS-use, verbal memory performance decline respectively. The differential changes between the HIGH and LOW group were located in the dorsal right striatum (putamen) (a). A higher putamen gray matter loss was associated with higher ATS-use and a higher cognitive decline between baseline and follow-up. In addition, higher interim ATS-use was associated with stronger cognitive decline (all p < 0.05).

## Discussion

Due to its prospective longitudinal design and the recruitment of occasional ATS users with very limited ATS exposure at study inclusion, the present study enabled a specific assessment of ATS use associated brain morphological changes. Importantly, users in the **LOW**and **HIGH**groups did not differ regarding previous or interim use of frequently co-used drugs, including cannabis and alcohol which often present severe confounders in the field and may affect striatal functional and structural integrity (4, 19) (Gouzoulis-Mayfrank et al., 2009; Zhou et al., 2019; Zimmermann et al., 2019; Gordin and Momenan, 2017). Together with the findings on a dose-response relationship in the correlational analyses, that took advantage of the entire sample, this suggests a direct association between the use of ATS and decreased dorsal striatal GM volumes and cognitive performance. Memory deficits, particularly immediate and delayed verbal memory, have been among the most consistently reported neurocognitive changes in ATS users, including chronic MA users (Scott et al., 2007; Roberts et al., 2018) as well as non-dependent populations such as recreational MDMA (Wagner et al., 2013; Schilt et al., 2007) and prescription AMPH users (Reske et al., 2010). However, despite the consistently observed functional impairments in the memory domain, and associated hippocampal memory functioning (Becker et al., 2013) evidence for altered structural hippocampal volume as a consequence of occasional or chronic ATS use is rather equivocal (Ersche et al., 2013; Daumann et al., 2011; Mackey et al., 2014; Berman et al., 2008). This may be may be related to the methodological properties of the VBM approach. Alterations in memory-related hippocampal functioning of ATS users are thought to due to changes in serotonergic (5HT) functioning (Wagner et al., 2013; Schilt et al., 2007). However, whereas previous studies that combined in-vivo receptor PET and VBM indicate a strong positive association between regional GM volume and dopaminergic D2/D3 receptor binding (Woodward et al., 2009), associations between 5HT receptor distribution and regional GM volume have not been reported (e.g. Jedema et al., 2010), suggesting that longitudinal VBM might have a higher sensitivity to detect alterations in DA pathways.

In line with the longitudinally observed associations between occasional ATS-use and GM changes, previous cross-sectional studies revealed some evidence for brain structural effects of occasional ATS use. A large study in occasional ATS and cocaine users revealed increased putamen and decreased inferior parietal GM volumes in occasional users as compared to non-using controls (Mackey & Paulus, 2014). In contrast, a previous cross-sectional comparison made by our group between baseline data from the occasional ATS users in the present study and drug-naïve controls did not reveal alterations in GM volume, probably due to the low-dose ATS exposure at study inclusion (Daumann et al., 2011). In addition, several cross-sectional studies examined brain morphological markers of chronic ATS use in MA-dependent individuals. Comprehensive reviews and meta-analytic evaluations of these cross-sectional comparisons revealed accumulating evidence for a consistent pattern of decreased prefrontal GM volumes accompanied by increased dorsal striatal, particularly putamen volumes in chronic MA users relative to controls (Ersche et al., 2013; Mackey & Paulus, 2014; Mackey et al., 2014). Several studies reported that within the group of MA users increased putamen volume was inversely associated with cognitive dysfunction (Chang et al., 2005; Jernigan et al., 2005; Jan et al., 2012), suggesting that increasing the GM volume of the striatum may be a compensatory response to initial neurotoxic effects.

In contrast to the consistently observed increases in putamen GM volume in chronic ATS users we found a decreased volume in continuing low-dose ones. Taken together with the association between GM and verbal memory decline this might suggest that compensatory responses have not yet occurred in the present sample. Interestingly, one study examined effects of short-term ATS-exposure on cognitive functioning and brain structural markers in children who were exposed to MA prenatally (36). In line with the present findings children with MA exposure demonstrated relative reductions in striatal, including putamen, volume and cognitive deficits in the domains of attention and memory. Notably, verbal memory deficits were specifically associated with the volume of the globus pallidus and the putamen (Chnag et al., 2004).

Studies examining the effects of ATS use at the molecular level have repeatedly observed deficient dorsal striatal DA neurotransmission in chronic MA users associated with functional deficits in motor and memory functioning (Volkow et al., 2001; Taylor et al., 2013). Likewise controlled studies in non-human primates observed decreased markers of dopaminergic functioning in the putamen following escalating MA regimes (Groman et al., 2013) as well as dopaminergic deficits in the dorsal striatum after low-dose AMPH exposure (Ricautre et al., 2005). Notably, an escalating MA regimen caused regionally-specific increased GM volumes in the putamen (Groman et al., 2013). Together with the previously reported correlation between regional GM volume and dopaminergic functioning (Woodward et al., 2009) this might suggest that the present findings parallel altered DA functioning in the dorsal striatum as a consequence of ATS use.

Although the dopaminergic basal ganglia (BG) system has been traditionally implicated in motor functioning and procedural learning (Bornelli & Cummings, 2007; Robbins et al., 2008) more recent evidence from BG disorders, particularly Parkinson disease (PD), lesion studies and pharmacological neuroimaging studies have revealed that the dorsal striatum contributes to learning and memory (Ward et al., 2013; Grahn et al., 2009). Cognitive impairments, most consistently in the domains of learning and memory, are a well-recognized feature in the early stages of PD (Grahn et al., 2009). Dopaminergic deficits in the putamen present the primary pathology during these initial stages of the disorder (Rodriguez-Oroz et al., 2009; Owen et al., 1998) and functional impairments show an extreme sensitivity to DA modulation (Lange et al., 1992), suggesting that they have a primary DA substrate. In addition, loss of putamen volumes has specifically been associated with cognitive deterioration in other neurodegenerative disorders characterized by marked memory impairments, including Alzheimer’s disease (de Jong et al., 2008). Moreover, evidence from patients with focal lesions to the BG, including the putamen, revealed impairments in the cognitive domains of working and verbal memory (Ward et al., 2013). One prospective study examining brain structure and cognitive functioning in 73 patients after carbon monoxide poisoning reported that stronger verbal memory impairments were associated with smaller putamen volumes 6 months following poisoning (Pulsipher et al., 2006). Recently, putamen volume has been genetically associated with schizophrenia, which is a psychiatric disorder characterized by its cognitive deficits (Luo et al., 2019). Although our knowledge of the role of the putamen in cognitive functioning is still incomplete these findings, together with the present results, indicate that alterations to its structure and function may result in more substantial cognitive impairment than previously assumed.

Altough the present prospective longitudinal design allowed to control for several important confounders inherent to retrospective design the findings have to be interpreted in the context of several limitations. First, several of the participants did not participate in the follow-up assessment and thus the sample size is comparably low and the findings need to be replicated in larger populations. Second, the study protocol and the target regions were not pre-registered and thus the findings should be considered as exploratory. Third, longer follow-up periods are necessary to determine the maintenance or recovery of the cognitive and brain functional changes over longer abstinence periods.

In summary, the present study has provided the first longitudinal evidence that prolonged use of low-dose ATS is associated with decreased dorsal striatal GM volume and verbal memory deficits, possible reflecting alterations in DA functioning.

## Author contribution

EGM, BB and JD were responsible for the study concept and design. DW and PK contributed to the acquisition of the data. BB, KMK and MT contributed to the analyses. BB and KMK interpreted the data and drafted the manuscript. All authors provided a critical revision of the manuscript, reviewed content and approved the final version for publication.

## Acknowledgements

K. Kendrick and J. Daumann contributed equally to this work (shared first authorship). This work was supported by the National Key Research and Development Program of China (Grant No. 2018YFA0701400). Q. Luo was supported by the National Key Research and Development Program of China (grant 2018YFC0910503), National Natural Science Foundation of China (grant 81873909), Shanghai Municipal Science and Technology Major Project (grant 2018SHZDZX01), Natural Science Foundation of Shanghai (grant 20ZR1404900), and Zhangjiang Lab. The authors thank all volunteers for their participation in this study. The authors declare that they have no conflict of interest.

## References

Aguilar MA, Garcia-Pardo MP, Parrot AC. Of mice and men on MDMA: A translational comparison of the neuropsychobiological effects of 3,4-methylenedioxymeth-amphetamine (“Ecstasy”). Brain Research. 2020;1727:146556

Ashburner J, Friston KJ. Diffeomorphic registration using geodesic shooting and Gauss-Newton optimisation. NeuroImage. 2011;55:954–967.

Ashburner J, Friston KJ. Voxel-based morphometry--the methods. NeuroImage. 2000;11:805–821.

Ashburner J, Ridgway GR. Symmetric diffeomorphic modeling of longitudinal structural MRI. Frontiers in neuroscience. 2012;6:197.

Becker B, Wagner D, Koester P, Bender K, Kabbasch C, Gouzoulis-Mayfrank E, Daumann J. Memory-related hippocampal functioning in ecstasy and amphetamine users: a prospective fMRI study. Psychopharmacology. 2013;225:923–934.

Becker B, Wagner D, Koester P, Tittgemeyer M, Mercer-Chalmers-Bender K, Hurlemann R, Zhang J, Gouzoulis-Mayfrank E, Kendrick KM, Daumann J. Smaller amygdala and medial prefrontal cortex predict escalating stimulant use. Brain: a journal of neurology. 2015;138:2074–2086.

Berman S, O’Neill J, Fears S, Bartzokis G, London ED. Abuse of amphetamines and structural abnormalities in the brain. Annals of the New York Academy of Sciences. 2008;1141:195–220.

Bonelli RM, Cummings JL. Frontal-subcortical circuitry and behavior. Dialogues in clinical neuroscience. 2007;9:141–151.

Chang L, Cloak C, Patterson K, Grob C, Miller EN, Ernst T. Enlarged striatum in abstinent methamphetamine abusers: a possible compensatory response. Biological psychiatry. 2005;57:967–974.

Chang L, Smith LM, LoPresti C, Yonekura ML, Kuo J, Walot I, Ernst T. Smaller subcortical volumes and cognitive deficits in children with prenatal methamphetamine exposure. Psychiatry research. 2004;132:95–106.

Daumann J, Koester P, Becker B, Wagner D, Imperati D, Gouzoulis-Mayfrank E, Tittgemeyer M. Medial prefrontal gray matter volume reductions in users of amphetamine-type stimulants revealed by combined tract-based spatial statistics and voxel-based morphometry. NeuroImage. 2011;54:794–801.

de Jong LW, van der Hiele K, Veer IM, Houwing JJ, Westendorp RG, Bollen EL, de Bruin PW, Middelkoop HA, van Buchem MA, van der Grond J. Strongly reduced volumes of putamen and thalamus in Alzheimer’s disease: an MRI study. Brain: a journal of neurology. 2008;131:3277–3285.

Eickhoff SB, Stephan KE, Mohlberg H, Grefkes C, Fink GR, Amunts K, Zilles K. A new SPM toolbox for combining probabilistic cytoarchitectonic maps and functional imaging data. NeuroImage. 2005;25:1325–1335.

Ersche KD, Jones PS, Williams GB, Turton AJ, Robbins TW, Bullmore ET. Abnormal brain structure implicated in stimulant drug addiction. Science. 2012;335:601–604.

Ersche KD, Williams GB, Robbins TW, Bullmore ET. Meta-analysis of structural brain abnormalities associated with stimulant drug dependence and neuroimaging of addiction vulnerability and resilience. Current opinion in neurobiology. 2013;23:615–624.

Everitt BJ, Robbins TW. Drug addiction: updating actions to habits to compulsion ten years on. Annual Reviews in Psychology. 2016: 67; 23–15

Gouzoulis-Mayfrank E, Daumann J. Neurotoxicity of drugs of abuse--the case of methylenedioxyamphetamines (MDMA, ecstasy), and amphetamines. Dialogues in clinical neuroscience. 2009;11:305–317.

Gouzoulis-Mayfrank E, Daumann J. The confounding problem of polydrug use in recreational ecstasy/MDMA users: a brief overview. Journal of psychopharmacology. 2006;20:188–193.

Grahn JA, Parkinson JA, Owen AM. The role of the basal ganglia in learning and memory: neuropsychological studies. Behavioural brain research. 2009;199:53–60.

Grodin EN, Momenan R. Decreased subcortical volumes in alcohol dependent individuals: effect of polysubstance use disorder. Addiction Biology. 2017: 22; 1426–1437.

Groman SM, Morales AM, Lee B, London ED, Jentsch JD. Methamphetamine-induced increases in putamen gray matter associate with inhibitory control. Psychopharmacology. 2013;229:527–538.

Jan RK, Lin JC, Miles SW, Kydd RR, Russell BR. Striatal volume increases in active methamphetamine-dependent individuals and correlation with cognitive performance. Brain sciences. 2012;2:553–572.

Jedema HP, Gianaros PJ, Greer PJ, Kerr DD, Liu S, Higley JD, Suomi SJ, Olsen AS, Porter JN, Lopresti BJ, Hariri AR, Bradberry CW. Cognitive impact of genetic variation of the serotonin transporter in primates is associated with differences in brain morphology rather than serotonin neurotransmission. Molecular psychiatry. 2010;15:512–522, 446.

Jernigan TL, Gamst AC, Archibald SL, Fennema-Notestine C, Mindt MR, Marcotte TD, Heaton RK, Ellis RJ, Grant I. Effects of methamphetamine dependence and HIV infection on cerebral morphology. The American journal of psychiatry. 2005;162:1461–1472.

Klugah-Brown B, Di X, Zweerings J, Mathiak K, Becker B, Biswal B. Common and separable neural alterations in substance use disorders: a coordinate-based meta-analysis of functional neuroimaging studies in humans. Human Brain Mapping. 2020: 4459–4477.

Lange KW, Robbins TW, Marsden CD, James M, Owen AM, Paul GM. L-dopa withdrawal in Parkinson’s disease selectively impairs cognitive performance in tests sensitive to frontal lobe dysfunction. Psychopharmacology. 1992;107:394–404.

Luo Q, Xhen Q, Wang W, Desrivieres S, Quinlan EB, Jia T, Macare C, Robert GH, Cui J, Guedj M, Palaniyappan L, Kherif F, Banaschewski T, Bokde ALW, Buechel C, Flor H, Frouin V, Garavan H, Gowland P, Heinz A, Ittermann B, Martinot JL, Artiges E, Paillere-Martinot ML, Nees F, Papadopoulus D, Poustka L, Froehner JH, Smolka MN, Walter H, Whelan R, Callicot J, Mattay VS, Pausova Z, Dartiges JF, Tzourio C, Crivello F, Berman K, Li F, Paus T, Weinberger, DR, Murray RM, Schumann G, Feng J, Imagen consortium. Association of a schizophrenia-risk nonsynonymous variant with putamen volume in adolescents: a voxel-wise genome-wide association study. JAMA Psychiatry. 2019: 435–445.

Mackey S, Paulus M. Are there volumetric brain differences associated with the use of cocaine and amphetamine-type stimulants? Neuroscience and biobehavioral reviews. 2013;37:300–316.

Mackey S, Stewart JL, Connolly CG, Tapert SF, Paulus MP. A voxel-based morphometry study of young occasional users of amphetamine-type stimulants and cocaine. Drug and alcohol dependence. 2014;135:104–111.

Maldjian JA, Laurienti PJ, Kraft RA, Burdette JH. An automated method for neuroanatomic and cytoarchitectonic atlas-based interrogation of fMRI data sets. NeuroImage. 2003;19:1233–1239.

Moratalla R, Khairnar A, Simola N, Granado, N, Garcia-Montes JR, Proceddu PF, Tizabi Y, Costa G, Morelli M. Progress in Neurobiology. 2017; 155: 149–170.

Owen AM, Doyon J, Dagher A, Sadikot A, Evans AC. Abnormal basal ganglia outflow in Parkinson’s disease identified with PET. Implications for higher cortical functions. Brain: a journal of neurology. 1998;121 (Pt 5):949–965.

Parrott AC. MDMA, serotonergic neurotoxicity, and the diverse functional deficits of recreational “Ecstasy” users. Neuroscience and Biobehavioral Reviews. 2013; 1466–1484.

Parrott AC. Why all stimulant drugs are damaging to recreational users: an empirical overview and psychobiological explanation. Human psychopharmacology. 2015;30:213–224.

Pulsipher DT, Hopkins RO, Weaver LK. Basal ganglia volumes following CO poisoning: A prospective longitudinal study. Undersea & hyperbaric medicine: journal of the Undersea and Hyperbaric Medical Society, Inc. 2006;33:245–256.

Reske M, Eidt CA, Delis DC, Paulus MP. Nondependent stimulant users of cocaine and prescription amphetamines show verbal learning and memory deficits. Biological psychiatry. 2010;68:762–769.

Ricaurte GA, Mechan AO, Yuan J, Hatzidimitriou G, Xie T, Mayne AH, McCann UD. Amphetamine treatment similar to that used in the treatment of adult attention-deficit/hyperactivity disorder damages dopaminergic nerve endings in the striatum of adult nonhuman primates. The Journal of pharmacology and experimental therapeutics. 2005;315:91–98.

Robbins TW, Ersche KD, Everitt BJ. Drug addiction and the memory systems of the brain. Annals of the New York Academy of Sciences. 2008;1141:1–21.

Roberts CA, Quednow BB, Montgomery C, Parrott AC. MDMA and brain activity during neurocognitive performance: An overview of neuroimaging studies with abstinent “Ecstasy” users. Neuroscience and Neurobehavioral Reviews. 2018: 470–482.

Rodriguez-Oroz MC, Jahanshahi M, Krack P, Litvan I, Macias R, Bezard E, Obeso JA. Initial clinical manifestations of Parkinson’s disease: features and pathophysiological mechanisms. The Lancet Neurology. 2009;8:1128–1139.

Rogers G, Elston J, Garside R, Roome C, Taylor R, Younger P, Zawada A, Somerville M. The harmful health effects of recreational ecstasy: a systematic review of observational evidence. Health technology assessment. 2009;13:iii–iv, ix-xii, 1-315.

Schilt T, de Win MM, Koeter M, Jager G, Korf DJ, van den Brink W, Schmand B. Cognition in novice ecstasy users with minimal exposure to other drugs: a prospective cohort study. Archives of general psychiatry. 2007;64:728–736.

Scott JC, Woods SP, Matt GE, Meyer RA, Heaton RK, Atkinson JH, Grant I. Neurocognitive effects of methamphetamine: a critical review and meta-analysis. Neuropsychology review. 2007;17:275–297.

Taylor SB, Lewis CR, Olive MF. The neurocircuitry of illicit psychostimulant addiction: acute and chronic effects in humans. Substance abuse and rehabilitation. 2013;4:29–43.

Thompson WK, Holland D, Alzheimer’s Disease Neuroimaging I. Bias in tensor based morphometry Stat-ROI measures may result in unrealistic power estimates. NeuroImage. 2011;57:1–4.

UNODC: UNO WDR, United Nations Office on drug and crimes world drug report, 2011 http://www.unodc.org/documents/data-and-analysis/WDR2011. 2011.

UNODC. UNO WDR, United Nations Office on drug and crimes world drug report, 2014 http://www.unodc.org/documents/data-and-analysis/WDR2014. 2014.

Volkow ND, Chang L, Wang GJ, Fowler JS, Leonido-Yee M, Franceschi D, Sedler MJ, Gatley SJ, Hitzemann R, Ding YS, Logan J, Wong C, Miller EN. Association of dopamine transporter reduction with psychomotor impairment in methamphetamine abusers. The American journal of psychiatry. 2001;158:377–382.

Wagner D, Becker B, Koester P, Gouzoulis-Mayfrank E, Daumann J. A prospective study of learning, memory, and executive function in new MDMA users. Addiction. 2013;108:136–145.

Ward P, Seri, Cavanna AE. Functional neuroanatomy and behavioural correlates of the basal ganglia: evidence from lesion studies. Behavioural neurology. 2013;26:219–223.

Woodward ND, Zald DH, Ding Z, Riccardi P, Ansari MS, Baldwin RM, Cowan RL, Li R, Kessler RM. Cerebral morphology and dopamine D2/D3 receptor distribution in humans: a combined [18F]fallypride and voxel-based morphometry study. NeuroImage. 2009;46:31–38.

Zhou X, Zimmermann K, Xin F, Zhao W, Derckx R, Sassmannshausen A, Scheele D, Hurlemann R, Weber B, Kendrick KM, Becker B. Cue-reactivity in the ventral striatum characterizes heavy cannabis use, whereas reactivity in the dorsal striatum mediates dependent use. Biological Psychiatry: Cognitive Neuroscience and Neuroimaging. 2019: 4:751–762.

Zimmermann K, Kendrick KM, Scheele D, Dau W, Banger M, Maier W, Weber B, Ma Y, Hurlemann R, Becker B. Altered reward processing in abstinent dependent cannabis users: social context matters. European Neuropsychopharmacology. 2019: 29; 356–364.

